# Experimental introgression to evaluate the impact of sex specific traits on *Drosophila melanogaster* incipient speciation

**DOI:** 10.1101/569699

**Authors:** Jérôme Cortot, Jean-Pierre Farine, Benjamin Houot, Claude Everaerts, Jean-François Ferveur

## Abstract

Sex specific traits are involved in speciation but it is difficult to determine whether their variation initiates or reinforces sexual isolation. In some insects, speciation depends of the rapid change of expression in desaturase genes coding for sex pheromones. Two closely related desaturase genes are involved in *Drosophila melanogaster* pheromonal communication: *desat1* affects both the production and the reception of sex pheromones while *desat2* is involved in their production in flies of Zimbabwe populations. There is a strong asymmetric sexual isolation between Zimbabwe populations and all other “Cosmopolitan” populations: Zimbabwe females rarely copulate with Cosmopolitan males whereas Zimbabwe males readily copulate with all females. All populations express *desat1* but only Zimbabwe strains show high *desat2* expression. To evaluate the impact of sex pheromones, female receptivity and *desat* expression on the incipient speciation process between Zimbabwe and Cosmopolitan populations, we introgressed the Zimbabwe genome into a Cosmopolitan genome labelled with the *white* mutation, using a multi-generation procedure. The association between these sex-specific traits was determined during the procedure. The production of pheromones was largely dissociated between the sexes. The copulation frequency (but not latency) was highly correlated with the female—but not with the male—principal pheromones. We finally obtained two stable *white* lines showing Zimbabwe-like sex pheromones, copulation discrimination and *desat* expression. Our study indicates that the variation of sex pheromones and of mating discrimination depend of distinct—yet overlapping—sets of genes in each sex suggesting that their cumulated effects participate to reinforce the speciation process.

## INTRODUCTION

The quality of sensory cues exchanged by sex partners is crucial with regard to sexual isolation and selection (DARWIN 1871a; DARWIN 1871b; ANDERSSON 1994; COYNE AND ORR 2004). Chemical signals emitted by conspecifics (pheromones) are used by many insects to assess the sex, species, population and reproductive status of their potential partner (HOWARD AND BLOMQUIST 2005). The intraspecific variation of pheromones (and of other sensory signals) can enhance the divergence between partly sexually isolated populations, this ultimately leading to distinct species (WYATT 2014). The mechanisms initiating and/or reinforcing speciation can either occur before copulation (altered mate discrimination; morphological alteration of genital parts; (EBERHARD 1993; ARNQVIST 1998) or after copulation (gametic or genomic incompatibility; (MAYR 1963; HURST AND POMIANKOWSKI 1991; PRESGRAVES *et al.*). However, the chronological involvement of these alterations and their relative contribution to sexual isolation remain difficult to determine together with the potential impact of novel sensory signals (WYATT 2014).

Pheromone natural variants have been discovered in several insect orders. In the European corn borer moth (*Ostrinia nubilalis*), some populations diverge both for the ratio between two pheromonal isomers (*E*) and (*Z*) of the 11-tetradecenyl acetate, and for the male behavioral response to the blends with different (*E*)/(*Z*) ratio (KLUN AND MAINI). Also, in some moths, the production of a variant (or novel) pheromone was caused by the reactivation of a silent desaturase gene (ROELOFS *et al.* 2002), or by a sex- and species-specific variation in the expression of this gene (LASSANCE *et al.* 2013). Since the apparition of the “novel” female pheromone was not directly linked to the gene(s) coding for the male behavioral preference for this pheromone (LÖFSTEDT *et al.* 1989), male preference may have pre-existed prior to the apparition of this novel female pheromone (BUTLIN AND RITCHIE 1989; ROELOFS *et al.* 2002). Indeed, desaturases are fast evolving genes related to speciation events in moths and in Drosophila species (FANG *et al.* 2009; SHIRANGI *et al.* 2009; XUE *et al.* 2012).

In the cosmopolitan species *Drosophila melanogaster*, most—but not all—populations show random mating (HENDERSON AND LAMBERT 1982; BEGUN AND AQUADRO 1993; KOROL *et al* 2000; HAERTY *et al.* 2002; YUKILEVICH AND TRUE 2008). Among the few known exceptions to this panmictic rule, a strong case of asymmetrical sexual isolation was reported in Zimbabwe populations (Z) where Z females rarely mate with cosmopolitan (M) males whereas Z males readily mate with all females (WU *et al.* 1995). A link between this incipient speciation case and the variation of expression in a desaturase gene *(desat2)* was proposed based on the presence of a functional *desat2* gene in Z, but not M strains (TAKAHASHI *et al.* 2001; FANG *et al.* 2002). However, the expression of *desat2* is not totally abolished in M strains, but seems to be “only” strongly repressed (MICHALAK *et al.* 2007). While the *desat2* gene is largely involved in the production of a variant female cuticular hydrocarbon (CH): 5,9-heptacosadiene (5,9HD; (COYNE *et al.* 1999; DALLERAC *et al.* 2000), its effect in the increased production of 5-tricosene (5T) in Z males remains unknown (GRILLET *et al.* 2012). While both Z females and males show high levels of C5-desaturated CHs, in several west-African strains only females—but not males—produce high levels of C5-desaturated CHs (5,9HD in Tai strain; (PECHINE *et al.* 1988; SUREAU AND FERVEUR 1999). The *desat1* gene, flanking *desat2*, is expressed in all *D. melanogaster* strains (Z and M) and codes for the production of C7-desaturated CHs in males (7-tricosene = 7T) and in females (7,11-heptacosadiene = 7,11HD; (JALLON 1984; WICKER-THOMAS *et al.* 1997; MARCILLAC *et al.* 2005a). Surprisingly, *desat1* is also involved in the discrimination of sex pheromones and in the emission of other yet unidentified mating cues (MARCILLAC *et al.* 2005b; BOUSQUET *et al.* 2012; BOUSQUET *et al.* 2016).

The variation of the female heptacosadiene ratio (7,11HD/5,9HD) has apparently no or very little effect on male copulation (FERVEUR *et al.* 1996; COYNE *et al.* 1999). Moreover, the experimental variation of the male tricosene ratio (7T/5T) only partly affects mating preference in Z females (GRILLET *et al.* 2012). This suggests that the asymmetrical sexual isolation between Z and M populations involves female perception of other male—non acoustic—sensory cues (COLEGRAVE *et al.* 2000; GRILLET *et al.* 2012; GRILLET *et al.* 2018). The hypothesis of sexual isolation based on multiple sensory signals and/or systems is somewhat supported by the finding that the divergence of mating preference between Z and M populations depend on a highly polygenic control (HOLLOCHER *et al.* 1997; TING *et al.* 2001; KAUER AND SCHLÖTTERER 2004). Also, among several genes showing mating-dependent variation of expression, *desat2* is up-regulated in Z flies, but down-regulated in M flies, after mating (MICHALAK *et al.* 2007). Post-copulatory isolation mechanisms inducing partial gametic incompatibilities were also detected between Z and M populations (ALIPAZ *et al.* 2001).

Since the relationship between (*i*) the variation of *desat2* expression, (*ii*) the production of cuticular sex pheromones in males, and (*iii*) the asymmetrical sexual isolation of Z populations remains unclear, we explored the genetic relationship between these traits in both sexes. To mimic a natural process, which may have taken many more generations, we carried out a multi-generations “backcross-selection” procedure in the laboratory to progressively introgress a Z genome into a M line carrying the *white* mutation (*w*). We measured, at different time points of this procedure, the relationship between these sex specific traits. This allowed us to evaluate their potential influence on the incipient speciation process observed between *D. melanogaster* populations. Our final goal consisted to obtain stable ZW lines associating Z-like sex pheromones, copulatory discrimination and *desat* expression with the *w* mutation. Such lines could be used to test the effect of transgenes associated with the *w*^+^ marker.

## MATERIAL & METHODS

### Flies and stocks

*Drosophila melanogaster* strains were raised in 15 ml glass vials containing 4 ml of yeast/cornmeal/agar medium and kept in a breeding room at 24.5±0.5°C with 65±5% humidity on a 12:12h light/dark cycle (subjective day from 8:00am to 8:00pm). M-type flies were transferred every two/three days (and Z-type flies every five to seven days) to avoid larval competition and to regularly provide abundant progeny for testing and breeding. All behavioral experiments were performed under similar conditions.

We used two M-type strains: Canton-S (Cs), an old-established strain widely used in fly laboratories, and the Di/w strain which was established (in 2011) after ten backcrosses between the Dijon 2000 (Di2) strain and the w^1118^ strain which carries the *white*^1118^ mutation (*w*; carried by the X chromosome and providing white eyes; (GREEN 1996) in a Cs genetic background.

We also used the Zimbabwe 30 strain (Z30; also noted Z6 in (GRILLET *et al.* 2012), a Z-type line which was collected in 1990 in the Wildlife Reserve of Sengwa in Zimbabwe (kindly provided by Profs Jerry Coyne, Chicago Univ. and Aya Takahashi, Tokyo Metropolitan Univ.). Both Z30 male and female flies produce relatively high level of variant desaturated cuticular hydrocarbon (CH) isomers.

All tested flies were screened under light CO_2_ anesthesia 0-4h after adult emergence and kept in fresh food vials for four or five5 days before testing. Males used in behavioral tests were held individually while females used in all the tests (behavior, CH analysis, *desat* gene expression, genetic crosses) were kept in groups of five to ten individuals.

### Genetic Procedure: backcrosses, selection and line establishment

We designed a genetic procedure including three backcrosses sessions (BC1, BC2, BC3) alternating with three “Analysis & Selection” sessions (A&S #1-3; Fig. 1). The procedure was initiated with a cross between Di/w females (M-type; *w*) and Z30 males (Z-type; throughout our ms, all crosses and pairs are indicated as “females x males”; we used this mating scheme since the reciprocal cross would have produced rare mating and fewer progeny). Our aim consisted to create and select lines associating the *w* mutation in an increasing proportion of Z genome (ZW lines). Each BC session (consisting of a 7-generations procedure; Fig. S1) was followed by a “A&S” session—lasting 3 to 4 generations—during which male and female pheromones and mating ability were measured in BC lines (successively producing ZW-BC1, ZW-BC2 and ZW-BC3 flies). More precisely, during each BC session, the female progeny resulting of the previous generation was (back)crossed either to ZW siblings (with *w* eyes) or to Z30 males (this mating scheme was chosen given that meiotic recombination only occurs in Drosophila females). The initial “Di/w x Z30” cross produced F1 flies with 100% red eyes females (*w*^+^/*w* heterozygous) and 100% white eyes males (*w* hemizygous). F1 x F1 crosses yielded about 50% F2 white eyes females (*w/w* homozygous) which were backcrossed to Z30 males. A similar mating scheme was used for the two following generations: F3 females (*w*^+^/*w*) were mated with F3 *w* males to produce F4 *w* females backrossed to Z30 males. This yielded F5 *w*^+^/*w* females and *w* males which were mated to statistically produce 50% F6 *w* flies. Small groups of F6 *w* sisters and brothers were intra-line mated to induce several inbred ZW-BC1 lines (only producing *w* flies) whose individuals were tested during the subsequent “A&S session #1”. Then, only ZW-BC1 lines showing the best Z-like characters were kept for the next BC session. A similar procedure was used to produce and select ZW-BC2 and then ZW-BC3 lines. To reduce the remaining genetic variability, pairs of flies from the ZW-BC3 lines—among the two lines showing the best Z-like characters (during the “A&S session #3”)—were mated to induce 20 isoparental sublines (IsoP).

**Figure 1.**
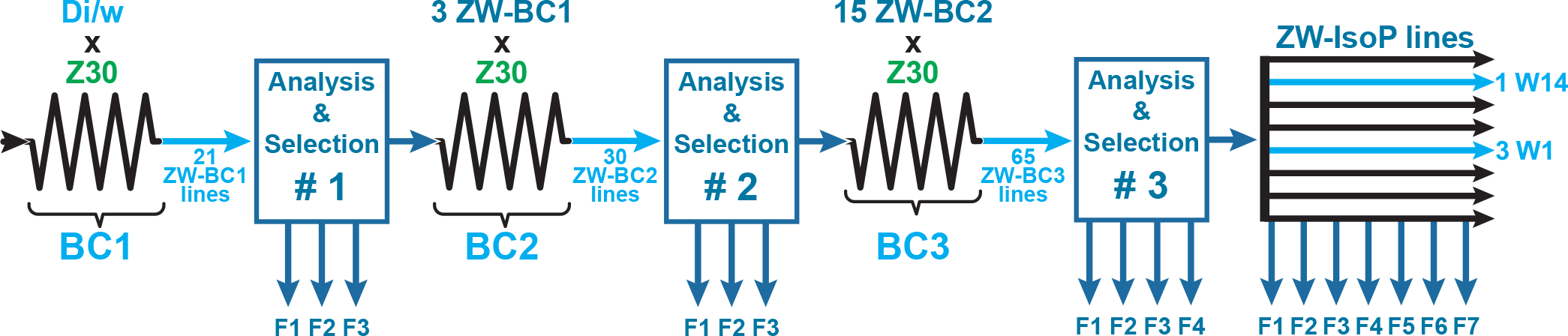
Genetic procedure of introgression. Dijon/w (Di/w) females were initially mated with Zimbabwe (Z30) males. During the following 6 generations, flies resulting of this cross were alternatively mated either between sisters and brothers, or females were backcrossed to Z30 males. The complete genetic procedure is detailed on Figure S1. From this “Backcross session #1” (BC1), we obtained 21 ZW-BC1 lines with mixed Z30 and Di/w genomes. Flies of these ZW-BC1 lines were analyzed for several phenotypes (“Analysis and Selection #1”) during the 3 following generations (F1, F2, F3), and the 3 lines showing the most Z30-like phenotypes were selected and kept to induce the next “Backcross session #2” (BC2). The BC2 session yielded 30 ZW-BC2 lines which were analyzed and selected (“Analysis and Selection #2”, during 3 generations: F1-F3). Among ZW-BC2 lines, 15 were kept to be involved in the BC3 session which yielded 65 ZW-BC3 lines which were analyzed and selected (“Analysis and Selection #3” during 4 generations: F1-F4). From this procedure, we isolated two lines (1W, 3W). From each line we created 20 isoparental lines (IsoP; one female x one male), which were analyzed at least during 7 generations (F1-F7). We finally retained two IsoP lines: 1W14 and 3W1.

### Mating behavior, fertility and fecundity

Behavioral tests took place 1-4h after lights on and were completed over several days for each genotype pair. Each male was individually aspirated (without an esthesia) under a watch glass used as an observation chamber (1.6 cm^3^) followed by a virgin female, 10 minutes later. Each test, performed under white light, lasted 120 minutes: for each pair, we noted the eventual latency of copulation (time lapse between the test start and copulation onset) and the overall frequency of copulating pairs for each treatment (for the sake of clarity, we do not show the copulation duration). In some experiments, when copulation occurred within the two-hour observation period, single mated females were individually placed into a food vial. Also, each non-mating pair (during the 2-hour test observation) was placed into a food vial and the male was discarded (with a mouth aspirator) after 24 hours. We only kept vials in which the mated female (in case of mating during the two-hour period) or the two flies of the pair were still alive 24 hours after the end of the mating test. In each vial, we determined both the fertility and fecundity based on the presence and number of adult progeny, respectively. The total number of viable adults yielded by each fertile pair was counted once, two weeks after the day of the mating test, and according to the mating status (mating or not during the 2-hours test).

### Cuticular hydrocarbons

5-Day old flies were frozen for 5 min at −20°C and individually extracted for 10 min at room temperature using 30 μl of hexane containing 3.33 ng/μl of *n*-C26 (*n*-hexacosane) and 3.33 ng/μl of *n*-C30 (*n*-triacontane) as internal standards. Male cuticular hydrocarbons (CHs) were quantified by gas chromatography using a Varian CP3380 gas chromatograph fitted with a flame ionization detector and an apolar CP Sil 5CB column (25 m by 0.25 mm; internal diameter: 0.1 μm film thickness; Agilent). In females, given that 5,9HD coeluted with 27-Br, we used a polar CP-Wax 58 FFAP column (25 m by 0.25 mm (internal diameter); 0.2 μm film thickness; Agilent). In both cases, a split–splitless injector (60 ml/min split-flow; valve opening 30 sec after injection) with helium as the carrier gas (50 cm/sec at 120°C) was used. The temperature program began at 120°C, ramping at 10°C/min to 140°C, then ramping at 2°C/min to 280°C and holding for 10 min. The chemical identity of the peaks was determined using a gas chromatography–mass spectrometry system equipped with a CP Sil 5CB or a polar CP-Wax 58 FFAP column as previously described (EVERAERTS *et al.* 2010). The amount (ng/insect) of each component was calculated based on the readings obtained from the internal standards. The overall sum of all the CHs (∑CHs) and of the principal CH groups were noted: desaturated CHs (∑Desat), linear saturated CHs (∑Lin) and branched CHs (∑Branched). We also show the ratio between ∑Desat and ∑Lin (D : L ratio). Ten to 20 flies were analyzed per sex. CH nomenclature is provided in Table S1.

Although the complete cuticular hydrocarbon profile were analyzed in all flies, for the sake of clarity, in most cases, we only show the predominant CHs (and their ratio) diverging between M and Z strains. This simplified CH analysis allowed us to screen in real time during successive generations, the CH profiles of many ZW-BC lines simultaneously sampled. However, the complete CH profiles of parental strains (Z30, Di/w, Cs) and of the two stable IsoP lines (1W14, 3W1) are shown (Table S1).

### Gene expression

To measure the relative amount of the five *desat1* transcripts (RA, RC, RE, RB and RD) and of the *desat2* transcript, we first extracted RNAs from 30 whole 5-day-old flies using the Trizol method (GIBCO BRL) and RNase-free DNase treatment to avoid contamination by genomic DNA (BOGART AND ANDREWS 2006). Total RNA (2μg) was reverse transcribed with the iScript cDNA Synthesis Kit (Biorad). Quantitative PCR reactions were performed with the IQ SYBR Green supermix (Biorad) in a thermal cycler (MyIQ, Biorad) according to the procedure recommended by the manufacturer. The qPCR reaction was done in a 20μl volume, by 40 cycles (95 °C for 30 sec, TM °C for 30 sec and 72 °C for 30 sec), preceded by a 3-min denaturation step at 98°C and followed by a 1-min elongation step at 72°C. TM of the hybridization step depends on the primer pair used (HOUOT *et al.* 2010). Each reaction was triplicated and the mean was calculated using three independent biological replicates. All results were normalized to the Actine5C mRNA level.

### Statistics

All statistical analyses were performed using XLSTAT 2012 (ADDINSOFT 2012). Frequencies were compared using a Wilks *G*^2^ likelihood ratio test completed with a computation of significance by cell (Fisher’s test). Comparison of fecundity, copulation latency and duration was carried out with a Kruskall-Wallis test with Conover-Iman multiple pairwise comparisons (*p*=0.05, with a Bonferroni correction). CH and/or behavioral data were analyzed using Principal Components Anaysis (PCA; type Pearson’s correlation matrix) with standardized values. For qPCR analysis, significant differences in transcript levels ratio between control and sample strain were detected with the Relative Expression Software Tool (REST, REST-MCS beta software version 2; PFAFFL 2001) where the iteration number was fixed at 2000. This test is based on the probability of an effect as large as that observed under the null hypothesis (no effect of the treatment), using a randomization test (Pair Wise Fixed Reallocation Randomisation Test©; PFAFFL *et al.* 2002).

## RESULTS

### 1) F0 and F1 flies phenotypes

Our genetic procedure consisted to progressively introgress the genome of a *D. melanogaster* wild-type strain collected in Zimbabwe (Z30 = Z) into the genome of a wild-type strain from Dijon (Di2) representative of M strains and carrying the *white* mutation (*w*; Di/w; Figs. 1 & S1; see Material and methods. We first measured the (*i*) cuticular pheromones and (*ii*) mating phenotypes in males and females of parental strains (F0 = Z30, Di/w, and Canton-S = Cs) and in the F1 progeny resulting of their reciprocal crosses (Z30 x Di/w; Z30 x Cs; Di/w x Z30; Cs x Z30; all crosses are shown as “females x males”).

For the sake of clarity, our pheromone analysis was mostly based on the measure of the 7T/5T ratio (T-ratio) in males and of the 7,11HD/5,9HD ratio (HD-ratio) in females. Both T-and HD-ratio showed much lower values in Z30 male and female flies (1.24±0.02 and 0.20±0.01, respectively) compared to M-type flies (for Cs: 14.85±0.34 and 31.31±0.63; for Di/w: 9.62±0.18 and 8.47±0.09, respectively). The absolute amounts (in ng) of these compounds, as well as the complete hydrocarbon profile of parental strains, are provided in supplemental information (Table S1)

Our mating experiments always involved pairs of flies (female x male; Fig. 2, Table S2). Since the *w* mutation carried by Di/w and ZW flies causes a visual defect inducing delayed mating latency, our behavioral observations lasted 2 hours. In parallel, and to control for a potential effect induced on mate discrimination by the *w* mutation, we also tested a second M-type strain with red eyes: the wild-type Canton-S strain (Cs). In the parental crosses, Z30 females showed, as expected, a contrasted mating frequency which was very low with both M males (4% with Cs; 0% with Di/w) and high with Z30 males (74%). Differently, both M females frequently mated with all males (60-98%), except in Cs x Di/w pairs (38%). F1 females frequently mated with Z30 males (69-85%) and with Cs males (59-83%). With Z30 and Cs females, most F1 males showed very high mating frequencies (73-100%) except for *w* F1 “Di/w x Z30” males with Z30 females (38%). This suggests that M-type genome induced a dominant effect over Z-type genome with regard to mating ability.

**Figure 2.**
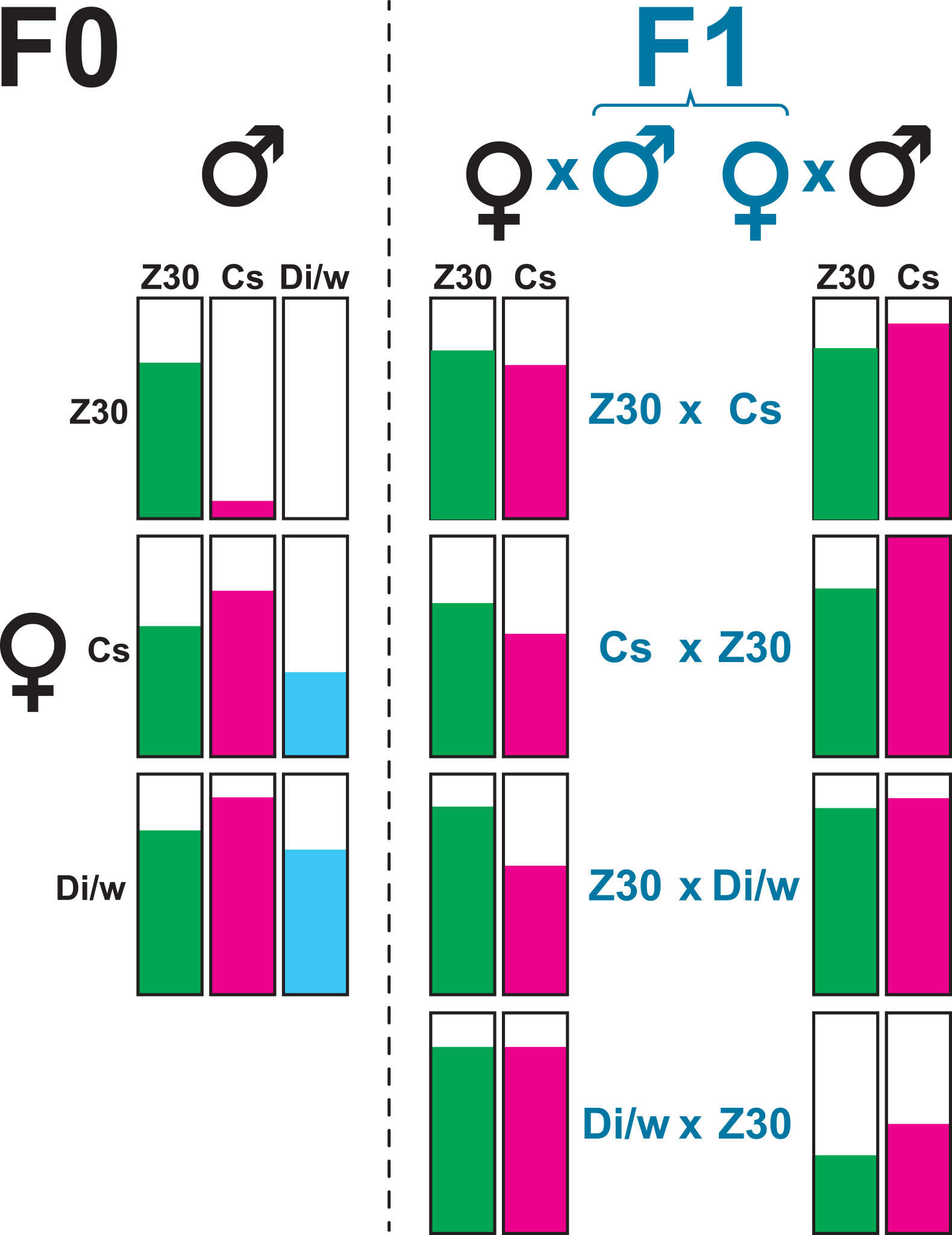
Copulation frequency in parental and F1 flies. We paired females and males from the three parental strains: Z30, Canton-S (Cs) and Dijon/w (Di/w; F0 left panel, indicated in black color). We also paired F0 flies with F1 flies resulting of the reciprocal crosses (indicated as: mother x father) between the three parental lines (F1 flies indicated in blue color). The filling within each bar represents the copulation relative frequency observed during two hours between pairs of flies.

Both the fertility (ability to leave progeny) and fecundity (number of adult progeny left) were measured in F0 pairs according to their copulation status (mating *vs.* not mating during the 2-hour test). While fertility and fecundity were not affected by copulation status (*r* range: −0.046 to 0.045; *p*=NS), both parameters were highly correlated in pairs mating within 2 hours or in the following 24h (*r*=0.689 and 0.828, respectively; *p*<0.05; Fig. S2). Z30 females showed a much higher fertility with Z30 males (74%) than with Cs and Di/w males (39 and 12%, respectively; G^2^ Wilks _(10df)_ =234.84) whereas M-type females always showed a very high fertility regardless of the male strain (88-98%). (Table S3). Moreover, Z30 x Z30 pairs copulating during the 2-hour period showed a much lower fecundity (median value=20 adults) compared to all other pairs involving M females (50-90 adults; KW_(6df)_=70.03; p<0.0001; Fig. S3).

### 2) Selection of ZW-BC1 lines

As mentioned above, the progeny of the “Di/w x Z30” cross was mated following the genetic procedure designed to establish ZW-BC1 lines with white eyed flies (*w*; see supra; Fig. S1 and Material and methods section).

We obtained 21 ZW-BC1 lines in which we measured during the “A&S session #1”: (1) male and female cuticular pheromones and (2) ZW male copulation with Z30 and Cs females. In particular, the T-ratio and HD-ratio were analyzed during 3 successive generations (F1, F2, F3; see “BC1” in Fig. 3). Given time limitation, we could only analyze one or two fly pool(s) per generation (pool size=3-11 for males; 2-6 for females).

**Figure 3.**
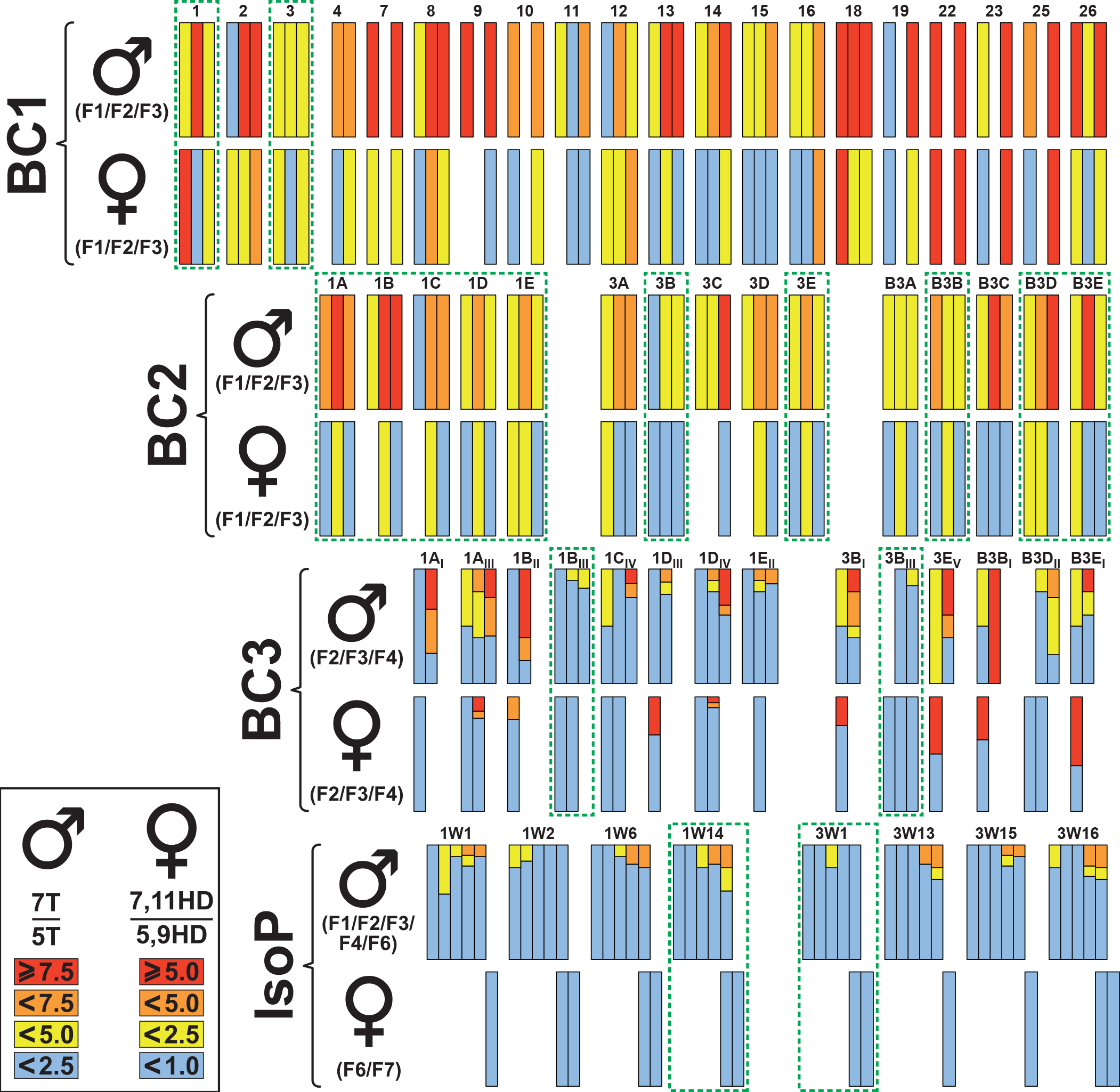
Cuticular hydrocarbons in male and female flies of selected lines during the introgression procedure. During each “Analysis and Selection session” we measured the ratio between 7-tricosene and 5-tricosene (7T/5T) in males and between 7,11-heptacosadiene and 5,9-heptacosadiene (7,11HD/5,9HD) in females. For the sake of clarity, we used a color code (from red to blue) corresponding to decreasing value of both ratio (see values in the inlet at the bottom left). More specifically, we measured these ratio in ZW-BC1 (BC1; F1-F3), ZW-BC2 (BC2; F1-F3), ZW-BC3 (BC3; F1-F3) and isoparental (IsoP; F1-F4, F6 in males, F6, F7 in females) lines. The number identifying each line is indicated above corresponding bars. The homogeneous color shown within each bar represents the mean value (calculated based on one or several pools of flies; BC1 and BC2) while the multiple colored bars (BC3, IsoP) represent the variability based on individual values. The green dotted frames indicate the lines selected and kept to the next step of the introgression procedure.

The T-ratio showed a wide variation in F1 (1.4-19.0), F2 (2.3-18.5) and F3 males (3.4-20.1). Similarly, the HD-ratio varied in F1 (0.1-47.9), F2 (0.1-4.9) and F3 females (0.2-21.1). Despite the presence of some extreme data points, the variability of both HD- and T-ratios tended to decrease between generations (Figs. S4 and S5A, respectively). A significant positive correlation between both T- and HD-ratio was found in F1 (*r*=−0.60; *p*=0.03), but not in F2 and F3 flies (*r*=0.48; *p*=0.10, and 0.29; *p*=0.21, respectively; Fig. S5B). This effect maybe partly explained by the negative correlation between absolute amounts of 7,11HD and 5,9HD which were higher in F1 (*r*=−0.77; *p*=0.002) than in F2 and F3 flies (*r*=−0.46; *p*=0.12, and −0.50; *p*=0.026, respectively). Similarly, but to a lesser extent, the amounts of C5- and C7-desaturated CHs showed higher level of correlation in F1 flies than in F2 and F3 flies (Fig. S6). The values for T- and HD-ratios are shown in Table S4.

While no F2 male copulated with Z30 females (within two hours), their mating frequency was highly variable with Cs females (between 9.8% for #12 line to 70% for #2 line; Table S5). However, given that these mating data were not enough discriminant between lines, we selected the three best ZW-BC1 lines only based on their CH phenotypes. Therefore, the #1, #3, #12 lines showing the lowest and most stable T- and HD-ratio values (especially in F3 flies) were used to initiate the BC session -2 (BC2).

### 3) Selection of ZW-BC2 lines

ZW-BC1 flies of #1, #3 and #12 lines reciprocally mated to Z30 parents yielded 6 ZW-BC2 main lines (#1, #3 and #12, resulting of Z30 x ZW-BC1 crosses; #B1, #bB3 and #B12, from ZW-BC1 x Z30 crosses). Each of these six main lines was subdivided into five sublines (labeled A to E) yielding a total of 30 ZW-BC2 sublines.

During the “A&S” session #2, both the CH profile and mating performance were analyzed in F1, F2 and/or F3 flies. After screening both CH (on pools) and mating phenotypes (see below) in F1 flies, we only kept the 15 sublines derived from the #1, #3 and B#3 ZW-BC2 lines (Figs. 3 & 4; the three other main lines, #B1; #12 and #B12, were further discarded). In most of the 15 sublines, the HD-ratio in F1 and F3 females was relatively low and stable (often<1.5; range=0.2-3.0), whereas the T-ratio showed a broader variability between and within sublines (for F1: 1.2-10.3; F2: 2.8-15.6; F3: 3.1-12.3; “BC2” in Fig. 3). The relatively high correlation value between T- and HD-ratios in F1 flies (*r*=0.60; *p*=0.03; Fig. S5B) strongly decreased in F2 flies (*r*=0.29; *p*=0.21). Such decrease maybe due to the low correlation between 7,11HD and 5,9HD and/or or null correlation between 7T and 5T, in F2 flies (Fig. S6). The values for T- and HD-ratios are shown in Table S6.

The mating performance of ZW-BC2 flies with M and Z30 flies substantially varied between lines (BC2 in Fig. 4; Table S7). At F1 generation, ZW x Cs pairs showed higher mating frequency in #1 (average=44.8%) than in #3 and #B3 sublines (3.7 and 7.0%, respectively). Note that females of the three other lines (further discarted: #B1; #12 and #B12) showed higher mating frequency with Cs males (9.7-23.1%). At F2 generation, ZW x Cs pairs showed a relatively homogeneous intra-line performance (#1=48.5% ranging from 35.0 to 61.1%; #3=6.2 —0 - 12.5%; #B3=15.9 —0 - 31.8%) compared to F1. Differently, Z30 x ZW pairs showed a broader intra-line variation: #1 (14.3%; 0-36.4%), #3 (25.4%; 9.1-56.6%) and #B3 (20.0%; 8.3-33.3%). At F3 generation, most Z30 x ZW pairs showed an increased mating frequency (#1=29.7% —25.0-33.3%; #3= 42.5% —33.3-56.3%; #B3=25.6% —12.5-43.8%). At F2 generation, a highly significant negative correlation was detected between the T-ratio of ZW males and their mating frequency with Z30 females (*r*=−0.71; *p*=0.004; Fig. S7A). Moreover, the copulation frequency in these pairs was negatively correlated with their copulation latency (Fig. S7B). However, no other significant correlation was found between any other CH-ratio and mating parameter in Z30 x ZW pairs (F2 and F3), or in ZW x Cs pairs (F1 and F2; Fig. S7). Based on these data, and as indicated above, we kept the 15 sublines derived from the #1, #3 and B#3 main ZW-BC2 lines to initiate the backross series #3 (BC3; Fig.1).

**Figure 4.**
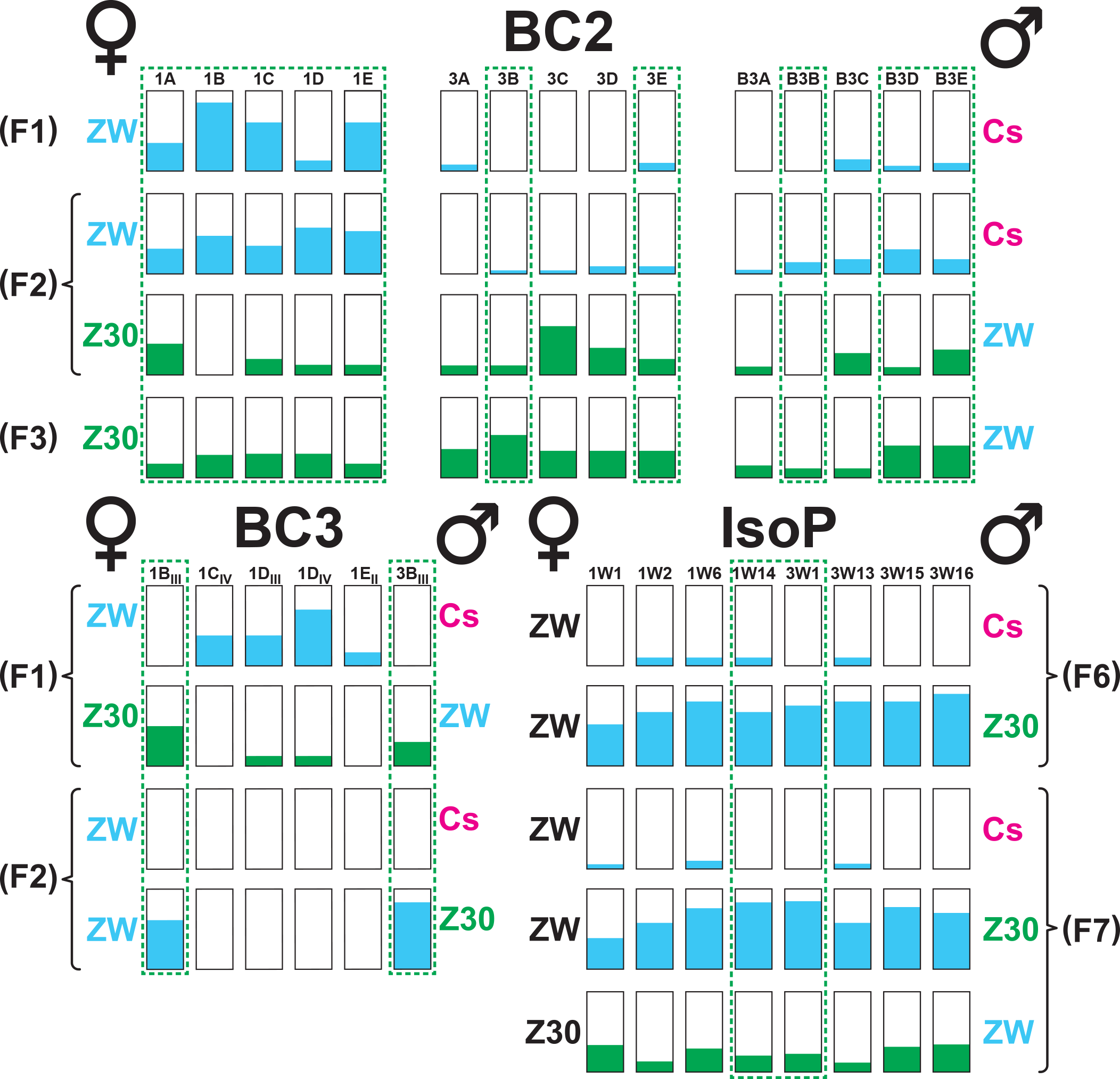
Copulation performance in flies selected during the introgression procedure. Each horizontal series of bars represents the copulation frequency (during two hours) of ZW-BC2 (BC2), ZW-BC3 (BC3) and isoparental (IsoP) flies reciprocally mated with parental Cs and Z30 flies. Experimental flies are indicated as “ZW”. For example, the top line represents ZW-BC2 females (at the F1 generation) paired to Cs males. The copulation relative frequency is indicated by the colored filling within each bar (see Figure 2). For all other information, please refer to Figures 1–3.

### 4) Selection of ZW-BC3 lines

Backcrossing ZW-BC2 females of the 15 selected sublines to Z30 males yielded 65 ZW-BC3 lines. More precisely, single females yielded by the last “BC3” generation were individually mated with 2-3 sibling males to initiate BC3-ZW isofemale sublines (labeled with I-VII roman numerals for each BC2 “mother” line). Each subline was first characterized by its T-ratio, given that this parameter seems to reflect best the proportion of “Z30” genome in ZW recombinant lines. The T-ratio was first measured (at F1 and F2) in two fly pools and subsequently—but only in lines showing a low T-ratio—characterized in individual males (at F3 and F4; “BC3” in Fig.3). When the four F1-F4 generations were considered, the T-ratio remained constantly low in only two sublines (#1B_III_ and #3B_III_; Table S8) while others produced at least some individuals (or pools) with higher T-ratio (≥5.0). The HD-ratio remained low in several ZW-BC3 sublines including #1B_III_ and #3B_III_, at least until F4 (≤0.3; Fig. 3; Table S9).

No significant “T-ratio/HD-ratio” correlation was found in F2 or F3 flies, despite a slight increase between these generations (*r*=0.04; *p*=0.89, *r*=0.25; *p*=0.43, respectively; Fig. S5B). This variation may be due to the increased correlation between the absolute amounts of 7,11HD and (*i*) 5,9HD or (*ii*) 5T (both negative) or (*iii*) 7T (positive) in F3 flies compared to F2 flies (Fig. S6).

We measured the mating performance of F1 and F2 flies showing the best Z30-like CH-ratio (#1B_III_, #1C_IV_, #1D_III_, #1D_IV_, #1E_II_ and #3B_III;_ “BC3” in Fig.4). At the F1 generation, 1B_III_ and #3B_III_ females did not copulate with Cs males (differently from the four other ZW-BC3 females), while 1B_III_ and #3B_III_ males substantially mated with Z30 females. At F2, #1B_III_ and #3B_III_ females showed a Z-like sexual discrimination: no copulation (within 2 hours) with Cs males and a high mating frequency with Z30 males (62 and 77%, respectively; Fig.4; Table S10). The mating frequency within and between the two ZW-BC3 lines was relatively low. While no correlation was detected in F3 ZW-BC3 x Cs pairs between any mating parameter and CH-ratio (Fig. S7A), a significant correlation was found between their copulation frequency and latency (Fig. S7B).

### 5) Selection of isoparental lines (IsoP)

To reduce as much as possible the potential intra-line genetic variability, pairs of ZW-BC3 flies were mated to initiate 20 “isoparental” lines derived from each #1B_III_ and #3B_III_ line (IsoP: 1W1-1W20 and 3W1-3W20, respectively). In each 1W and 3W line, the “F1 to F4” stability of the T-ratio was assessed using individual males (“IsoP” in Fig. 3; Table S11). Based on these data, only four 1W isoparental sublines (#1, 2, 6 and 14) and four 3W isoparental sublines (#1, 13, 15 and 16) were kept for subsequent testing. In these 8 IsoP sublines, F6 and/or F7 females showed a low HD-ratio (≤0.3; Table S12) whereas the T-ratio diverged between lines. Based on these data, we only retained the “1W14” and “3W1” lines which showed a stable T-ratio until F9. A great similarity for the complete CH profiles was found between 1W14, 3W1 flies and parental Z30 flies, in both sexes (Table S1). Our most recent analysis carried out with F75 flies show a very stable profile (data not shown)

Simultaneous testing of the 8 IsoP lines (F6-F7) revealed a weak correlation between T- and HD-ratio (*r*=0.12, p=0.79) caused by the opposite correlation between 1W and 3W lines (respectively, *r*=−0.65 — *p*=0.35 and *r*=0.54 —*p*=0.46) (Fig. S5B). The correlation between the absolute amounts of single CHs was relatively low and not significant between 5T and either 7,11HD (*r*=0.33; *p*=0.42) or 5,9HD (*r*=0.35; *p*=0.40; Fig. S6). However, a very high correlation was found within the 3W1 line (but not the 1W14 line) between 7- and 5T (*r*=0.64; *p*=0.01) and between 7,11- and 5,9HD (*r*= 0.71; *p*=0.002).

F6 and F7 IsoP females showed a high mating discrimination: they rarely mated with Cs males (0-10%) and much more frequently with Z30 males (44-90%; “IsoP” in Fig. 4). F7 IsoP males moderately mated with Z30 females (11-36%), or with F7 IsoP sibling females (6-28%; Table S13).

We performed a further cross-examination of CH profile with mating performance in IsoP lines using Principal Component Analysis (PCA; Fig. S9A). This revealed that while (*i*) both T- and HD-ratios were not correlated together (*r*=0.026), (*ii*) the copulation frequency of Z30 x IsoP pair was correlated with HD-ratio (*r*=0.524), (*iii*) the copulation frequency of both IsoP x IsoP and IsoP x Cs pairs was correlated with the T-ratio (*r*= 0.420 and 0.414, respectively), but (*iv*) the copulation frequency of IsoP x Z30 was not correlated with any CH ratio (*r*=− 0.041 and −0.075).

We also determined the fertility and fecundity of F6- and F7 IsoP flies used in mating tests. IsoP females showed higher fertility with Z30 males (65-92%) than with Cs males (16-32%; Table S14) whereas IsoP male fertility was high with both Z30 or IsoP females (71-85% and 78-100%, respectively). This suggests that mating events occuring after the two-hours observation period were more frequent in IsoP x IsoP and IsoP x Z30 pairs than in IsoP x Cs pairs.

The fecundity, measured with regard to the genotypes and mating status of pairs (Fig. S8) only revealed slight difference. IsoP x Z30 and Z30 x IsoP fertile pairs left around 20 and 30 adult progeny per cross, respectively with no difference between IsoP lines. A slight effect of mating status was detected in IsoP x Z30 pairs: copulating pairs showed a slightly higher fecundity than non-copulating pairs (U=3588, *p*=0.037). IsoP x IsoP fertile pairs generally produced a low progeny number (<20) but no effect was detected between lines.

### 6) *desat* gene expression in IsoP lines

The transcriptomic profiles of the *desat1* and *desat2* genes were compared between the three F0 lines and the 1W14 and 3W1 lines (sampled at F8 and F9 generations). More precisely, we measured the level of the five *desat1* transcripts (RA, RC, RE, RB and RD) and the unique *desat2* transcript, in mature flies of both sexes. Males and females of the Z-type lines (Z30, 1W14 and 3W1) showed significantly higher levels of the *desat2* transcript compared to both M-type parental strains (Cs; Di/w shown as the baseline of Fig. 5). The *desat1* transcripts showed much less interstrain difference. In both sexes, the three Z-type lines showed a similar—although reduced—variation compared to M-type strains. The only notable exception was detected for the RE transcript which decreased (*i*) in Z-type females compared to M-type females and (*ii*) in Cs males compared to the four other males.

**Figure 5.**
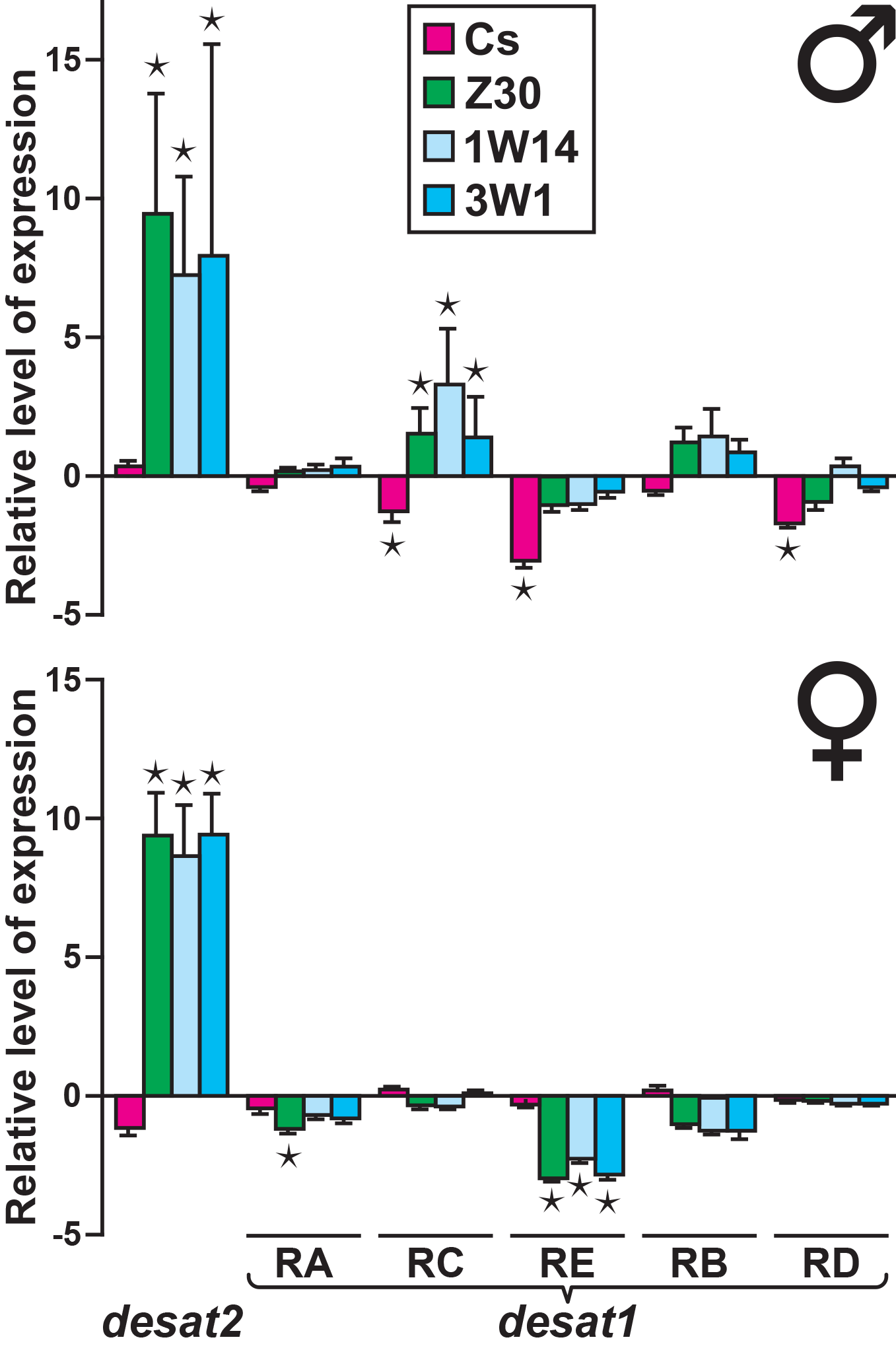
Relative levels of *desat1* and *desat2* transcripts expression in parental and isoparental introgressed lines. Each set of bars indicates the relative positive or negative variation of the single *desat2* transcript and each of the five *desat1* transcripts (RA, RC, RE, RB, RD) in male (top) and female flies (bottom). The baseline axis corresponds to the values measured in Di/w flies of respective sex. Colored bars indicate the relative variation in parental Cs and Z30 lines and in the two isoparental lines 1W14 and 3W1. Statistical differences between the values of Di/w and the four other lines are indicated (*: *p*<0.05). *N*=3 biological replicates.

## DISCUSSION

The experimental introgression of a Z genome into a *white*-labeled M genome allowed us to follow, during many generations, the evolution of the relationship between several sex specific traits diverging between *D. melanogaster* populations. In particular, we found that cuticular pheromones and female sexual receptivity—two sex specific traits potentially involved in incipient speciation—depend of distinct sets of genes. After more than 20 generations of introgression and 10 generations of experimental selection, we obtained two white-eyes lines (1W14 and 3W1) showing the principal Z-type characters: female sexual discrimination, male copulation with Z30 female, male and female desaturated CHs, fecundity and *desat1*-*desat2* gene expression.

Based on our data, the chronological scenario of the events involved in this specific case of incipient speciation remains hypothetical. Together with previous reports, our data indicate that Z- and M-type strains diverge for several aspects involved in both pre- and postzygotic isolation. Postzygotic isolation is revealed by the lower fertility and fecundity found in Z30 and IsoP pairs. This is supported by other studies (ALIPAZ *et al.* 2001). The prezygotic isolation is reflected by the positive correlation between mating frequency and HD-ratio. More specifically, high 7,11HD level was positively correlated with mating frequency while increased (thus delayed) copulation latency was correlated with high levels of 5,9HD (Fig. 6). However, while increased levels of the two HD isomers induced reciprocal effect on both copulation parameters, the HD-ratio variation was only correlated with the frequency of copulation but not with its latency (Fig. S9B). This apparent conundrum suggests that the behavioral effect induced by both HD isomers does not only depend on their absolute amounts *per se* but also on their relative contribution to the pheromonal bouquet, this determining their ratio. Other studies have shown that the ratio between two compounds with potential pheromonal effect, or their relative contribution to a more complex bouquet, can strongly influence insect ability to discriminate between closely related individuals, colonies, sub-species or species (KLUN AND MAINI 1979; COLLINS AND CARDÉ 1985; ADAMS AND HOLT 1987; OGUMA *et al.* 1992; FERVEUR AND SUREAU 1996; AYASSE *et al.* 1999; MARCILLAC *et al.* 2005c; LIN *et al.* 2010; VAN ZWEDEN AND D’ETTORRE 2010). Given that the correlation between the absolute amounts of the two HD isomers tended to decrease during the introgression procedure (except in BC3-F3; Fig. S6), this suggests that the production of these compounds does not only depend of *desat2*, which may nevertheless exerts a major influence on the HD-ratio (COYNE *et al.* 1999).

**Figure 6.**
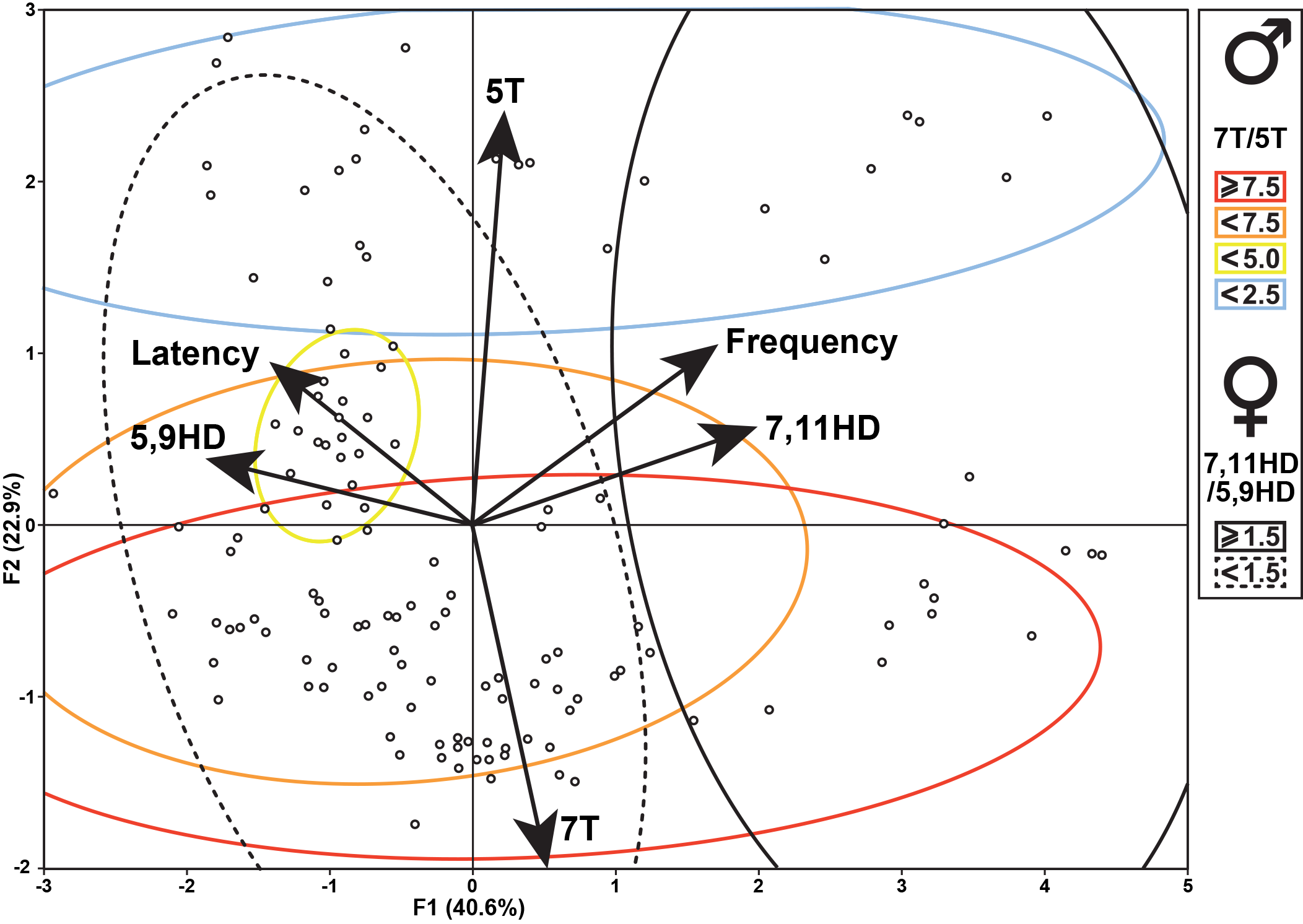
Relationship between the principal cuticular hydrocarbons and copulation parameters during the introgression procedure. When ZW-BC1, ZW-BC2 and ZW-BC3 flies (but not IsoP) were considered, we found a high correlation between (*i*) the frequency of copulation success rate with the 7,11HD level and (*ii*) the copulation latency with the 5,9HD level. No relationship was found between the copulation latency and frequency, or between 5T, or 7T and either copulation parameter. The amount of variability for the two principal axes is indicated (F1: 40.6%; F2: 22.9%). The distributions of male and female cuticular hydrocarbons ratio are included in ellipses represented with four colors (males) and by plain/dotted lines (in females; see inlet shown on the right).

Differently, no clear relationship was detected between the mating ability and either the absolute amounts of two principal tricosene isomers (7T, 5T; Fig. 6), or their ratio (T-ratio; Fig. S9B). This may be due to the fact that our global statistical analysis including all ZW-BC sessions masked more subtle effects. For instance, during the introgression procedure the sign of the correlation between the absolute amounts of 7T and 5T changed: it was negative in F1 ZW-BC1 (*r*=−0.40) and became positive in IsoP lines (*r*=0.25; Fig. S6). Despite the absence of a global significant correlation, our genetic procedure largely based on the selection of “low T-ratio” in ZW males allowed us to isolate two stable IsoP lines combining the *w* mutation with all main Z-like sex specific phenotypes. This suggests that some of the genes (with Z-type alleles) involved in the low T-ratio and in mate discrimination are closely linked. This suggests that these hypothetic “low T-ratio” coding genes could also affect non-pheromonal Z male traits (sensory signals, behavioral postures) together with traits underlying female receptivity to Z male traits.

Several *D. melanogaster* studies revealed the involvement of single genes in female sexual discrimination and receptivity (SUZUKI *et al.* 1997; CARHAN *et al.* 2005; DITCH *et al.* 2005; JUNI AND YAMAMOTO 2009; SAKAI *et al.* 2009). This indicates that accurate female discrimination of male characters depends of many genes organized in networks (GREENSPAN 2001; FERVEUR 2005; HOUOT *et al.* 2012). Therefore, any alteration in each of these genes could induce detrimental effect on female mating behavior. Most of these genes may determine the production or the reception of each of the multiple sensory signals reciprocally exchanged during the courtship interplay by the two partners (MARKOW 1987; ARIENTI AND JALLON 1991; WELBERGEN *et al.* 1992; LASBLEIZ *et al.* 2006; KRSTIC *et al.* 2009). Taken together, these observations indicate that Z30 female discrimination depends of multiple genes shaping her accurate ability to perceive and integrate, during different phases of courtship, the multiple male sensory signals. Based on this, we believe that the origin of assortative mating between Z- and M-type females is linked to their divergent perception and/or integration of male cues. Indeed, while Z females only respond to a precise and complete multisensory set of signal provided by homotypic males (GRILLET *et al.* 2012; GRILLET *et al.* 2018), M-type females only need part of these cues to be sexually receptive (COLEGRAVE *et al.* 2000; GRILLET 2009; MA *et al.* 2010; GRILLET *et al.* 2012; GRILLET *et al.* 2018).

Our hypothesis that genes underlying the production of cuticular pheromones and mating responses at least partly differ, is also supported by the divergent dominance status of their Z- and M-type alleles: they both semi-dominantly control the production of C5- and C7- desaturated CHs (Fig. 2; Table S1) while M-type alleles dominantly act over Z-type alleles with regard to female discrimination (Fig. 2). Our data also reveal that the production of C5- and C7-desaturated CHs depends on a distinct genetic determination between the sexes: a stable Z-like HD-ratio was obtained after a relatively low number of generations in ZW-BC females while it took many more generations to stabilize a Z-like T-ratio in ZW-BC males. As a consequence, the variation between HD- and T-ratios was not correlated through the introgression process. This indicates that the production of C5- and C7-desaturated CHs at least partly depends of a sex specific control: the HD-ratio mostly—but not only (see above)—depends of the *desat2* gene (COYNE *et al.* 1999), while the T-ratio may additionally depends on many other genes. Some of the “low T-ratio coding genes” diverging between Z and M populations and potentially involved in sexual isolation maybe influenced by mating activity (HOLLOCHER *et al.* 1997; TING *et al.* 2001; KAUER AND SCHLÖTTERER 2004; MICHALAK *et al.* 2007). Indeed, two genes detected in the later study (CG12400; *Cyp4p2*) code for enzymes potentially involved in hydrocarbon biosynthesis (JALLON 1984; QIU *et al.* 2012). The hypothesis of a sex specific control for the production of C5- and C7-desaturated CHs isomers is also supported by the dissociation of their production between the sexes in West-African strains (Tai strain females predominantly produce 5,9HD while Tai males produce 7T and 7P; PECHINE *et al.* 1985; FERVEUR *et al.* 1996; SUREAU AND FERVEUR 1999).

We still do not understand how Z strains have been able to survive in nature given all their potentially disadvantageous reproduction-related characters compared to M strains which have spread all over our planet. The Z-type populations may be very strictly adapted to local ecological conditions only found in Zimbabwe forests. Based on their genetic diversity and the simultaneous expression of both *desat1* and *desat2* genes, Zimbabwe populations have been proposed to represent ancestral *D. melanogaster* populations from which M-type populations have derived. Then, during their global expansion on earth (maybe in relation with human migration; DAVID AND CAPY 1988), M populations may have progressively adapted their chemosensory perception system together with other non-chemosensory perception systems (ARGUELLO *et al.* 2016).

In conclusion, the present study offers a “real-time measure” of the evolution of several sex-specific traits potentially involved in sexual isolation. Our data provide a statistic view on their relative involvement and also on their inter-sex relationship. We are now planning to introduce transgenes targeting chemosensory genes (including *desat1*) into the genome of our two stable ZW lines to further dissect the mechanisms underlying the incipient speciation process between *D. melanogaster* populations to better understand the recent evolution and worldwide adaptation of this species.

## Acknowlegments

This work was partly funded supported by the Centre National de la Recherche Scientifique (INSB), the Burgundy Regional Council (PARI 2012), the Université de Bourgogne, the ANR (Gustaile) and the CONICYT (MEC 80140013).

---

**Figure S1. Detailed genetic procedure for the first backcross session**. Dijon/w (Di/w) females were initially mated with Zimbabwe (Z30; green color) males. Resulting F1 ZW females (with colored eyes; magenta color) were mated with resulting F1 ZW males (white eyes; cyan color). Only resulting F2 ZW females with white eyes (cyan color) were backcrossed to Z30 males (Bc: F2 x F0). Then, a “sister x brother” cross was carried on the next generation (F3 x F3) between white-eyed flies. A backcross between F4 white-eyed females and Z30 males (F4 x F0), was followed by two successive “sister x brother” crosses (F5 x F5; F6 x F6) to yield sublines with homogeneous white-eyed progeny. Flies resulting of the F6 x F6 “sister x brother” cross produced white-eyed ZW-BC1 females and males which were tested in the “Analysis and Selection #1” session (see Figure 1).

**Figure S2. Relationship between copulation status, fertility and fecundity in F0 strains**. Using the 9 possible crosses between the 3 F0 parental lines (Cs, Di/w, Z30; see Figure 2), we compared the fertility (ability to leave at least one adult progeny) and the fecundity (number of adult progeny per female) in females copulating within the two-hour observation period (2H) or during the following 24-hour period (>2H). The PCA reveals a high correlation between fertility and fecundity for each copulation status but not between copulation status. The amount of variation taken into consideration for each axis is indicated (F1: 42.26%; F2: 30.69%).

**Figure S3. Fecundity in crosses between parental flies according to copulation status**. The number of adult progeny per female was counted in each of the cross involving a male (upper row under the bars) and a female (bottom row) according to the copulation status: mating within the two-hours observation period (COPULATION 2H) or during the 24 hours period (COPULATION >2H). Box-plots represent the 50% median data (the small horizontal bar indicates the median value, and the solid dot represents the mean; the whiskers represent the first and third quartiles). When the sample sizes are too small, single data points are shown as crosses. Different italic lowercase letters above whiskers indicate significant differences between means.

**Figure S4. Variation of cuticular hydrocarbons ratio during the introgression procedure**. For each graph, the two dimensions axes represent the absolute amount of cuticular hydrocarbons (in ng). For the x-axis: 7, 11HD in females (left panels) and 7T (in males, right panels); For the y-axis 5,9HD and of 5T, respectively. The distribution for these two pairs of compounds (in females and in males) is shown for parental (F0) Z30 (green ellipse) and Di/w (cyan) flies. The dotted lines indicate the ratio between the two compounds for F0 flies together with three additional “intermediate” ratio values. The values shown for BC1, BC2 and BC3 (Analysis and Selection) sessions are shown according to the generation at which they were sampled. (See Figure 3).

**Figure S5. Relationship between female and male cuticular hydrocarbons ratio**. (**A**) The distribution of the female (7,11HD/5,9HD) and male (7T/5T) ratio is shown on a two axis diagram according to the generation sampled during the introgression procedure. The position of the F0 Di/w line is highlighted by an arrow on the right upper corner. (**B**) Bars represent the level of the correlation between the 7T/5T ratio and the 7,11HD/5,9HD ratio at different generations of the introgression process. The significant correlations are indicated (*: *p*<0.05).

**Figure S6. Correlation between cuticular hydrocarbons during the introgression procedure**. The level and sign of the correlation between pairs of CHs (indicated under the corresponding bars) are shown at different generations of the introgression procedure (the color code for generations is shown on the right side).

**Figure S7. Correlation between copulation parameters and cuticular hydrocarbons ratio**. Bars indicate the level and the sign of the correlation between (**A**) the 7/5 ratio (7,11HD/5,9HD in ZW females; 7T/5T in ZW males) and the copulation frequency (dark bars) or the copulation latency (light bars) or (**B**) the copulation frequency and latency. The sex and generation of the pairs of flies tested are indicated under each set of histogram bars. Iso 1W and Iso 3W correspond to four pooled 1W IsoP and four pooled 3W IsoP lines, respectively. The statistical significance is indicated near each corresponding bar.

**Figure S8. Fecundity in isoparental lines**. The number of adult progeny left by each pair was counted according to its mating status (“2H”: copulation during two-hour period; “>2H”: copulation during the following 24-hour period:) and the origin of each parent: females are shown on the top row (under box-plot bars) and males on the bottom row. Flies of isoparental lines (IsoP; four 1W and four 3W IsoP lines) were reciprocally crossed with Z30 flies; IsoP females were also paired with Cs males and with IsoP males of the same line. For more information, please refer to Figure S3).

**Figure S9. Relationship between cuticular hydrocarbons and copulation performance**. (**A**) The PCA represents the relationship between copulation frequency in females and/or males of eight selected isoparental lines (IsoP) paired with parental flies (Cs, Z30) together with CH ratios (7,11HD/5,9HD in females; 7T/5T in males). A high correlation was found between the 7T/5T ratio and the copulation frequency in IsoP x Cs (IsoP females x Cs males) and with IsoP pairs. Another high correlation was detected between the 7,11HD/5,9HD ratio and the copulation frequency of Z30 females paired with IsoP males (Z30 x IsoP). The relative variability is indicated for the two principal axes (F1: 32.30%; F2: 22.97%). (**B**) When ZW-BC1, ZW-BC2 and ZW-BC3 (not IsoP) flies were considered, a very high correlation was found between the frequency of copulation success rate and the 7,11HD/5,9HD ratio. No correlation was found between these two parameters and the copulation latency or with the 7T/5T ratio; the two latter parameters were not correlated together. The amount of variability for the two principal axis is indicated (F1: 44.2%; F2: 30.7%). The distributions of male and female cuticular hydrocarbons ratio are included in ellipses represented with four colors (males) and by plain/dotted lines (in females; see inlet on the right side).

